# Heart Disease in a Mutant Mouse Model of Spontaneous Eosinophilic Myocarditis Maps to Three Highly Significant Loci

**DOI:** 10.1101/747519

**Authors:** Nives Zimmermann, William J. Gibbons, Shelli M. Homan, Daniel R. Prows

## Abstract

**Background:** Heart disease (HD) is the major cause of morbidity and mortality in patients with hypereosinophilic diseases. Due to a lack of adequate animal models, our understanding of the pathophysiology of eosinophil-mediated diseases with heart complications is limited. We have discovered a mouse mutant, now maintained on an A/J inbred background, that spontaneously develops hypereosinophilia in multiple organs. Cellular infiltration into the heart causes an eosinophilic myocarditis, with affected mice of the mutant line (*i.e.,* A/J^HD^) demonstrating extensive myocardial damage and remodeling that leads to HD and premature death, usually by 15-weeks old. Maintaining the A/J^HD^ line for many generations established that the HD trait was heritable and implied the mode of inheritance was not too complex. Backcross and intercross populations generated from mating A/J^HD^ males with females from four different inbred strains produced recombinant populations with highly variable rates of affected offspring, ranging from none in C57BL/6J intercrosses, to a few mice with HD using 129S1/SvImJ intercrosses and C57BL/6J backcrosses, but nearly 8% of intercrosses and >17% of backcrosses from SJL/J related populations developed HD. Linkage analyses of these SJL/J derived recombinants identified three highly significant loci: a recessive locus mapping to distal chromosome 5 (LOD=4.88; named *Emhd1* for <u>e</u>osinophilic <u>m</u>yocarditis to <u>h</u>eart <u>d</u>isease-<u>1</u>); and two dominant variants mapping to chromosome 17, one (*Emhd2*; LOD=7.51) proximal to the major histocompatibility complex, and a second (*Emhd3*; LOD=6.89) that includes the major histocompatibility region. Haplotype analysis identified the specific crossovers that defined the *Emhd1* (2.65Mb), *Emhd2* (8.46Mb) and *Emhd3* (14.59Mb) intervals. These results indicate the HD trait in this mutant mouse model of eosinophilic myocarditis is oligogenic with reduced penetrance, due to multiple segregating variants and possibly additional genetic or nongenetic factors. The A/J^HD^ mouse model represents a unique and valuable tool to understand the interplay of causal factors that underlie the pathology of this newly discovered eosinophil-associated disease with cardiac complications.

## Introduction

Discovering a spontaneous mouse mutant and the genetic variants underpinning its disease can be invaluable in the pursuit of understanding the corresponding human disease. Once identified, these mutants can provide the means to investigate the disease, understand the associated pathological mechanisms and discover new therapies. Usually, natural mutants involve a single *de novo* gene mutation that often can be identified quickly with current mapping and next generation sequencing technologies. However, the causal variants for about half of spontaneous mutants have evaded discovery by exome sequencing [1]. Many reasons can explain why mutations go undetected using exome sequencing, including poor gene annotations, poor exon capture, intronic mutations or variants in upstream or downstream regulatory regions, structural and copy number variants, epigenetic changes, and environmental effectors. In addition, it is likely that many unmapped mutant traits result from decreased penetrance due to multiple segregating genetic variants. One explanation for the sudden discovery of a complex trait is that one or more contributing variants lie dormant and do not cause disease on their own. But, when the effects from a critical new mutation combines with the prior latent variant(s), the mutant trait is revealed. In this setting, this oligogenic trait would appear to be Mendelian.

In this report, we describe the genetic mapping of a complex trait discovered in our mouse colony. From the founder spontaneous mutant, we have established a novel mouse line on a SNP-verified A/J inbred background. Affected mice of this line naturally (*i.e.,* uninduced) develop an eosinophilic myocarditis (EM) that progresses to heart disease (HD) and death, with almost three-quarters of affected mice dying by 15-weeks old [2]. Penetrance of the disease trait is reduced and varies greatly among the breeder pairs, consistent with multiple genetic variants still segregating for the EM/HD trait. However, we have maintained this mutant line (designated as A/J^HD^) for many years without a reliable screening marker, suggesting that heritability of the trait is not exceedingly complex. Litter ratios of affected-to-unaffected offspring show a large and breeder-pair specific range. However, extensive pedigree data for hundreds of matings over many generations implicates 2–3 genes (depending on their modes of inheritance) segregating in the A/J^HD^ mutant line, supporting that mapping the EM/HD trait should be feasible.

Using quantitative trait locus (QTL) analysis, we have discovered three highly significant loci, designated *Emhd1–3* for ‘<u>e</u>osinophilic <u>m</u>yocarditis to <u>h</u>eart <u>d</u>isease’. These three QTLs include a homozygous recessive variant residing on distal chromosome 5 (*Emhd1*) and two distinct QTLs on chromosome 17; one (*Emhd2*) that maps proximal to the major histocompatibility complex (MHC) and a second interval (*Emhd3*) that includes the MHC. Evaluating the mice used for QTL mapping, haplotype analysis identified specific recombinations that delineated the proximal and distal ends of each QTL interval. Together, these three QTLs explain ∼64% of total trait variance, implicating the potential involvement of one or more other factors. The established A/J^HD^ line offers a powerful resource to identify the set of causal variants (coding and noncoding) underlying these three QTLs and any additional genetic or non-genetic factors that contribute to the initiation or progression of disease. Knowing the genes involved in EM/HD will allow targeted studies to understand the pathologic mechanisms at play for this and possibly other eosinophil-related diseases with cardiac manifestations, for which little genetic information is currently recognized.

## Results

### General pathology of A/J^HD^ mutant mice

Whole-mount images of hearts from an A/J^HD^ litter are presented in **Fig. 1**. No indications of HD were observed in three siblings (Fig. 1A-C), whereas signs of HD were noted in three of their littermates, including one heart of approximately normal size with significant atrial and ventricular fibrosis (Fig. 1D) and two with extensive right ventricular dilatation and dramatic right outer wall thinning that approached transparency (Fig. 1E-F). To confirm the extensive fibrosis seen in the myocardium of affected mutants, hearts from mice with HD were stained with Trichrome. A representative affected mutant heart **(Fig. 2B**) with images taken at several sites (indicated by arrows) and increased magnifications (Fig. 2C-H) and compared to an unaffected A/J^HD^ littermate (Fig. 2A).

**Fig. 1.**
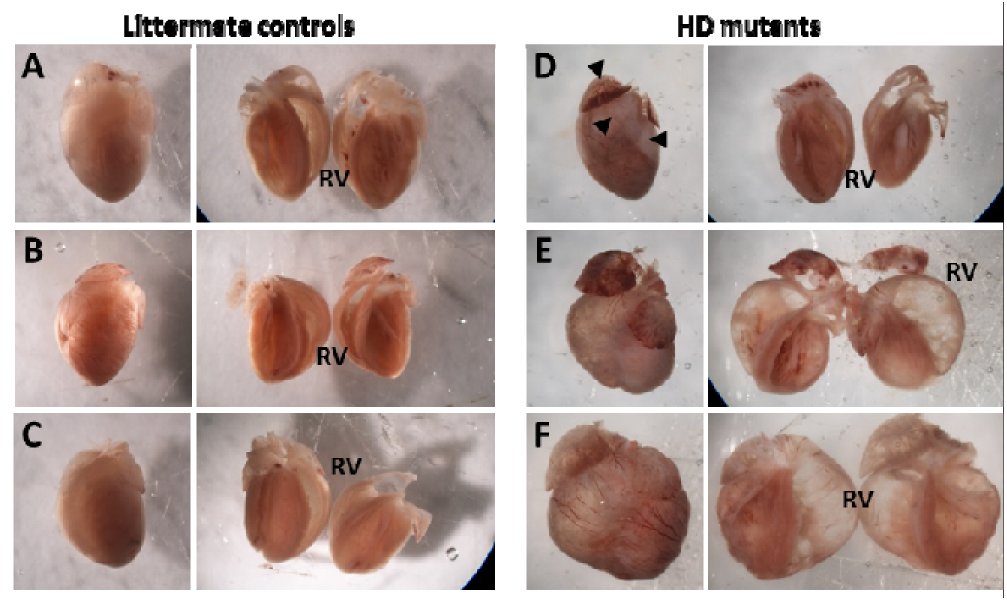
Dissection microscope photos. Whole mounts with coronal cuts for hearts removed from littermate controls (**A–C**) and three mutants (**D–F**). All three mutants had visible signs of heart disease. Two mutants (**E**, **F**) show evidence of left atrial enlargement, right ventricular dilation with dramatic right wall thinning, and left ventricle hypertrophy. A third mutant (**D**) presents as slightly smaller than controls, but with significant atrial and ventricular fibrosis (arrow heads). All photos were taken at the same settings.

**Fig. 2.**
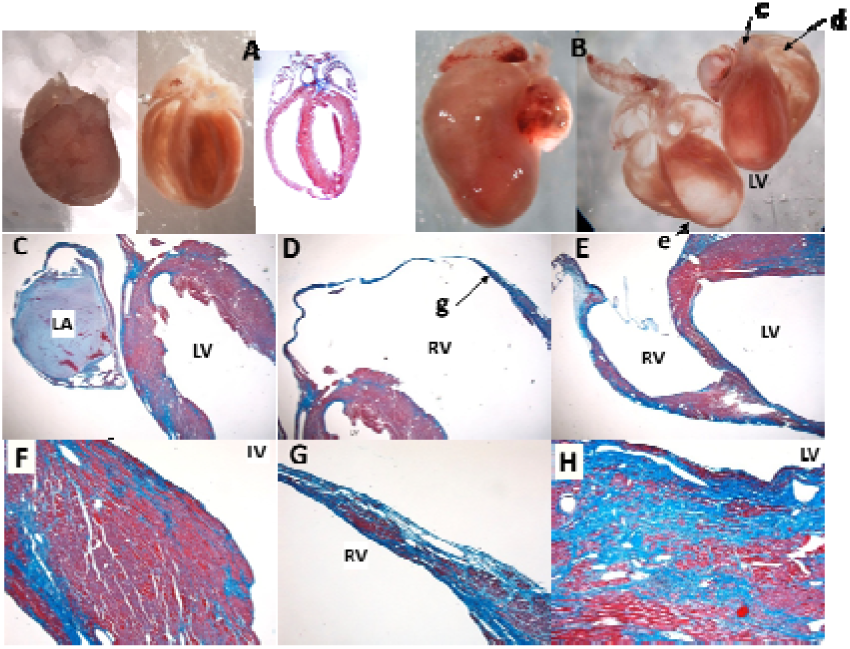
Whole mounts and Trichrome staining of hearts. Mutant hearts were coronally cut, paraffin mounted and stained with Trichrome. Unaffected control (**A**) shows little to no Trichrome staining, whereas the affected mutant heart (**B-H**) demonstrates extensive fibrosis in both ventricles. Lowercase letters (**c to g**) indicate the focal point of the corresponding magnified views on a light microscope (**C-F**).

### Replicating the trait in recombinants

We have sustained the mutant line for many years by maintaining about 10 breeder pairs that consist of brother-sister or parent-offspring matings and by using siblings of mice that had died of HD to help preserve the necessary pool of disease-associated alleles. To map these causal variants, our goal was to reproduce the disease trait dozens of times in recombinant mice generated from at least two different strains. To detect both dominant and recessive linkage, we first used an F_2_ breeding scheme. Initially, F_1_ hybrids produced from C57BL/6J (B6) females and proven A/J^HD^ mutant males (*i.e.,* males that had previously sired offspring developing HD) were intercrossed to generate a cohort of 102 B6.A/J^HD^ (B6.HD) F_2_ mice (Additional file 1, see 1A); unexpectedly, none of these B6.HD-F_2_ recombinants developed HD (Table 1). A second B6.HD-F_2_ population (Additional file 1, see 1B) was generated using different mutant male breeders, but also yielded no mice that developed HD (n=181; Table 1). Compared to the 2-3 genes estimated to be segregating in the A/J^HD^ inbred line, these B6.HD-F_2_ data (0 affected in 283 F_2_ mice) suggested a considerably more complex pattern of trait inheritance (*i.e.*, at least 5 variants or other genetic or non-genetic factors).

**Table 1.** Crosses used to produce mice for EM/HD mapping.

| Recombinant Cross | # litters | ♂, affected | ♀, affected | total, affected | % affected |
| --- | --- | --- | --- | --- | --- |
| (B6 x A/J<sup>HD</sup>) F<sub>2</sub> | 10 | 54, 0 | 48, 0 | 102, 0 | 0 |
|  | 15 | 84, 0 | 97, 0 | 181, 0 | 0 |
| (B6 x A/J<sup>HD</sup>) N<sub>2</sub> | 5 | 21, 0 | 29, 1 | 50, 1 | 2.0 |
| (S1 x A/J<sup>HD</sup>) F<sub>2</sub> | 32 | 123, 0 | 143, 2 | 266, 2 | 0.75 |
| (D2 x A/J<sup>HD</sup>) F<sub>2</sub> | 58 | 262, 9 | 248, 6 | 510, 15 | 2.9 |
| (C5 deficient) |  |  |  |  |  |
| (D2 x A/J<sup>HD</sup>) N<sub>2</sub> | 26 | 114, 7 | 122, 13 | 236, 20 | 8.5 |
| (C5 deficient) |  |  |  |  |  |
| (SJ x A/J<sup>HD</sup>) F<sub>2</sub> | 31 | 148, 12 | 139, 10 | 287, 22 | 7.7 |
| (Dysf deficient) |  |  |  |  |  |
| (SJ x A/J<sup>HD</sup>) N<sub>2</sub> | 13 | 42, 8 | 56, 9 | 98, 17 | 17.3 |
| (Dysf deficient) |  |  |  |  |  |

Our second attempt concurrently generated two distinct recombinant populations to further explore whether the HD trait could be recapitulated in a mapping cross (Additional file 1, see right). For these crosses, we changed the mating scheme for one recombinant population and switched the inbred strain for the second. The first population again used B6 females, but this time in a backcross strategy to potentially double our chances of recapitulating the trait (Additional file 1, see 2A). In this case, a small cohort of B6 backcrosses (B6.HD-N_2_) was produced by using each mutant A/J^HD^ male (proven sires) for both the initial F_1_ outcross and the subsequent backcrosses with his daughters; still, just one of 50 (2%) B6.HD-N_2_ recombinants developed HD (Table 1). Concomitantly, we generated 266 F_2_ recombinants from mutant A/J^HD^ males and 129S1/SvImJ (S1) inbred females (Additional file 1, see 2B); two S1.HD-F_2_ mice developed HD (0.75% affected). Thus, although rare, HD was recapitulated in two different inbred strain-pairs and two breeding schemes, demonstrating that mapping the trait might be possible; however, this would require a huge population of recombinant mice to produce the number of affected mice required for linkage analysis. Still puzzling, however, was the discrepant rates of HD seen between these mapping crosses and the inbred A/J^HD^ line.

### Improving the rate of affected recombinant mice

To explain the disparity between the estimated number of segregating causal mutations in the A/J^HD^ line compared to the recombinants from B6.HD-N_2_, B6.HD-F_2_, and S1.HD-F_2_ crosses, we hypothesized that one or more causal variants contributing to EM/HD must already exist as fixed polymorphism(s) in the A/J inbred strain (and the A/J^HD^ inbred mutant line). Because one or more of these A/J polymorphisms are fixed as homozygous, they are masked when maintaining the A/J^HD^ line with brother-sister and parent-offspring matings. However, these fixed mutations begin to segregate when the A/J^HD^ line is outcrossed with another inbred strain for the mapping studies, which significantly reduces the odds to reestablish the full set of allelic variants needed to cause HD. Given the large difference in the rates of affected offspring between A/J^HD^ and its mapping crosses, it is plausible that more than one latent A/J variant is affecting the HD trait. By choosing a strain with the same or functionally similar mutation as in A/J, we could indirectly test this possibility while still attempting to generate recombinants for mapping. Because of their potential role in EM/HD pathology, hemolytic complement (*C5*) and dysferlin (*Dysf*) were prioritized from the list of 14 known mutated disease genes in A/J inbred mice (see: https://www.jax.org/strain/000646; View Genetics).

#### C5

Two conceivable mechanisms of disease initiation include 1) an unknown pathogen acts as a trigger or 2) an unknown self-protein induces an autoimmune response. C5 has a potential role in both: C5 deficiency increases susceptibility of A/J mice to many infectious agents [3–6] and C5 function and dysfunction are indicated in immunity and autoimmunity [7–11]. From this, we predicted that the effects from the natural *C5* mutation in the A/J strain [12] combines with that of the *de novo* mutation (and possibly other latent mutations) in our A/J^HD^ line to cause disease. To test this hypothesis A/J^HD^ males were mated with DBA/2J (D2) females, an inbred strain that carries the same 2-bp deletion in *C5* as does A/J [13]. These matings necessarily produce litters in which all recombinant offspring have homozygous mutant copies of *C5* (as do the A/J and A/J^HD^ lines). If *C5* deficiency contributes to disease, then the odds are improved that the breeders will carry the required set of mutant alleles and produce more affected recombinant offspring.

D2 mice were bred using F_2_ and N_2_ mating schemes (Additional file 1, bottom) to evaluate a potential role of *C5* in EM/HD and to produce recombinants for mapping the causal variants of HD. Cohorts of 510 intercross mice (D2.HD-F_2_) and 236 backcross mice (D2.HD-N_2_) for a total of 746 D2-derived recombinants, were produced and monitored for HD up to 20-weeks old (Table 1). D2.HD-F_2_ recombinants developed HD at a rate of 2.9% (15/510), with no sex difference identified (*p*=0.186). Twenty D2.HD-N_2_ mice (8.5%) developed HD, with a 1.7-fold difference (*p*=0.017) between females (13/122; 10.7%) and males (7/112; 6.3%). As expected, because the same mutant male was used for the F_1_ and then the N_2_ crosses with daughters, the rates of affected D2.HD-N_2_ mice were 2.9-fold that of D2.HD-F_2_ mice (8.5% versus 2.9%). The use of D2 mice increased the percentages of affected recombinants over B6 and S1 mice, consistent with our prediction that the lack of C5 has a direct or indirect role in EM/HD, possibly stemming from a more susceptible immune milieu in the absence of C5.

#### Dysf

For an additional strategy to improve the rate of generating recombinants developing HD and to identify regions significantly linked to the disease, we tested a parallel hypothesis that the known *Dysf* gene mutation in the A/J strain (and thus the A/J^HD^ line) contributes to the mutant phenotype. Although the *Dysf* mutation in the A/J strain appears to be unique to this strain (a 5-6kb ETn retrotransposon inserted into intron 4) [13], the SJL/J (SJ) inbred strain has a splice-site mutation in *Dysf*, resulting in a markedly decreased protein level [14]. Like the A/J strain, the SJ strain is also considered a naturally occurring animal model for dysferlinopathy [15]. Consequently, all mice deriving from crosses of the SJ and A/J^HD^ strains will carry two mutated copies of the *Dysf* gene. Like the D2 crosses for mutant *C5*, an increase in affected SJ-derived recombinants would indirectly support an involvement of mutant *Dysf* in the development, progression, and/or modification of the pathobiology associated with EM/HD.

Accordingly, the SJ-derived crosses were used to map the causal HD genes and test a potential contributing role for *Dysf* in EM/HD; again, the N_2_ and F_2_ mating schemes were exploited (Additional file 1, left). Cohorts of 287 intercross mice (SJ.HD-F_2_) and 98 backcross mice (SJ.HD-N_2_), for a total of 385 SJ-derived recombinants were produced and monitored up to 20-weeks old for HD (Table 1). SJ.HD-F_2_ recombinants developed HD at a rate of 7.7% (22/287), with no sex difference observed (females: 10/139; 7.2% and males 12/148; 8.1%; *p*=0.348). Seventeen SJ.HD-N_2_ mice (17.3%) developed HD, with a slight yet significant (*p*=0.029) difference between females (9/56; 16.1%) and males (8/42; 19.0%). Again, the backcross breeding scheme produced HD at more than twice the rate of F_2_ crosses (17.3% versus 7.7%). Consistent with our hypothesis of a contributing role for *Dysf* deficiency in EM/HD, the use of SJ-derived crosses dramatically increased the percentages of affected recombinants. Comparing these final crosses to the first 3 attempts (Additional file 1), the SJ.HD-N_2_ (17/98; 17.3%) improved the rate of HD ∼35-fold over the combined B6 and S1 crosses (3/599; 0.5%).

### QTL analysis

Assessment of genetic linkage to the EM/HD trait was performed using R/qtl analysis [16, 17] to associate HD (binary trait: present or not by 20-weeks old) with SNPs on the GigaMUGA SNP array generated for all affected and unaffected mice in the analyses. Each cross was analyzed separately, then combined with its alternative mating scheme partner (*i.e.,* D2 or SJ crosses).

#### D2 crosses

For the D2.HD-N_2_ analysis, a total of 19 affected (*i.e.*, verified HD) and 6 unaffected (*i.e.,* littermates that did not develop HD by 20-weeks old) backcross mice were used. Similarly, 16 affected and 16 unaffected D2.HD-F_2_ mice were utilized in a separate QTL analysis (Table 2). Except for the chromosome 2 area around *C5*, the F_2_ population will help to rule out genomic regions that are homozygous D2 in affected mice.

**Table 2.** QTLs identified in R/qtl analyses.

| Cross<br>(#HD / #Unaffected) | Chr position<br>(@ peak SNP) | LOD<br>Score | % Variance<br>Explained | Level of<br>Significance* | Haplotype-defined interval<br>Chr: Mb range (size of interval) |
| --- | --- | --- | --- | --- | --- |
| D2.HD-N<sub>2</sub><br>(19 / 6) | 5:147147276 | 3.32 | 43.2 | suggestive | 5:143159090–151734385<br>(8.58 Mb) |
| D2.HD-F<sub>2</sub><br>(16 / 16) | – | – | – | – | 5:144329217–147894899<br>(3.57 Mb) |
| SJ.HD-N<sub>2</sub><br>(16 / 17) | 5:148558183 | 4.88 | 47.4 | highly significant<br>(*Emhd1*) | 5:129525220–151734385<br>(22.21 Mb) |
| SJ.HD-F<sub>2</sub><br>(22 / 22) | – | – | – | – | 5:145128608–147778059<br>(2.65 Mb) |
|  | 17:21456192 | 7.51 | 52.9 | highly significant<br>(*Emhd2*) | 17:15358323–23816187<br>(8.46 Mb) |
|  | 17:35308100 | 6.89 | 49.6 | highly significant<br>(*Emhd3*) | 17:31471362–45658739<br>(14.19 Mb) |
| SJ.HD-N<sub>2</sub> + F<sub>2</sub><br>(38 / 39) | 1:176044015 | 4.12 | 21.3 | significant | 1:172969406–194625219<br>(23.59 Mb) |
|  | 5:146601763 | 5.79 | 28.6 | highly significant<br>(*Emhd1*) | 5:143159090–151734385<br>(8.58 Mb) |
|  | 17:17933928 | 6.53 | 35.1 | highly significant<br>(*Emhd2*) | same as F<sub>2</sub> |
|  | 17:35308100 | 7.43 |  | highly significant<br>(*Emhd3*) | same as F<sub>2</sub> |

All genomic positions are based on mm10 (GRCm38; Build 38). Genotyping of >143,00 SNPs was performed by GeneSeek (Lincoln, NE) using the GigaMUGA SNP panel. Separate QTL analyses were run for each cross using R/qtl [16, 17] after applying argyle routines for quality checks [18]. Combined SJ crosses were analyzed as an F<sub>2</sub> model (0,1,2), accepting the limitations. \* Significance was determined using 1,000 permutations of the corresponding dataset. QTL intervals were determined using haplotype analysis, in which we identified the proximal and distal crossovers that best correlated the genotype with phenotype.

Results of the D2.HD-N_2_ QTL analysis identified a suggestive linkage on distal chromosome 5 (LOD=3.32; **Fig. 3**). Details of the peak markers, LOD scores, percent variance explained and QTL intervals for this and all other performed analyses are summarized in Table 2. Initially, the results of the D2.HD-N_2_ cohort also indicated a suggestive locus on distal chromosome 17 (LOD=3.39; Fig. 3), but this peak was subsequently ruled out after determining the region coincided with residual B6 genome from the founder A.B6 congenic. Specifically, one of the A/J^HD^ breeder males used in the D2.HD-N_2_ crosses still contained a portion of the original B6 congenic interval. However, since HD occurred in mutant mice after removing the residual B6 interval, linkage to this region (and thus the QTL peak) was ruled out. Interestingly, neither the D2.HD-F_2_ analysis nor a combined (D2.HD-N_2_ + D2.HD-F_2_) analysis identified any locus that reached suggestive linkage (Additional file 2).

**Fig. 3.**
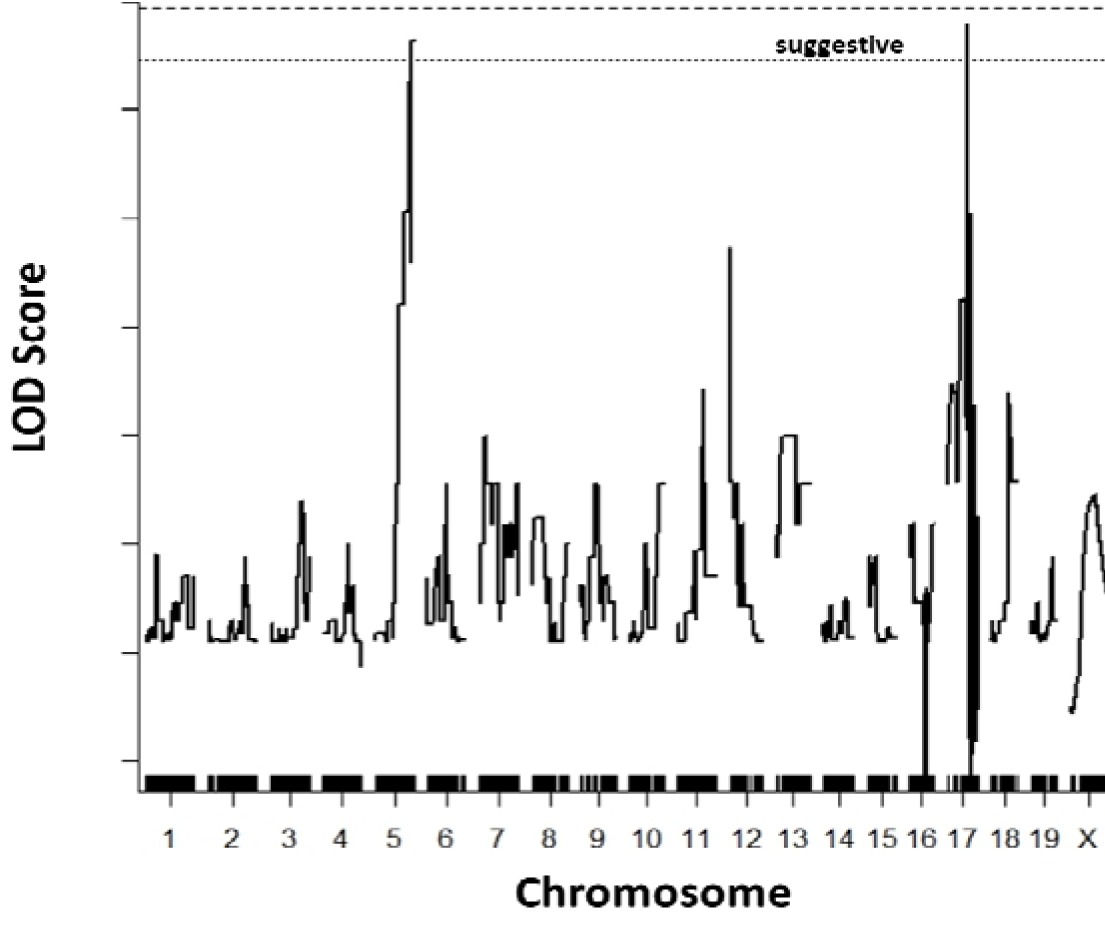
QTL analysis results of the D2.HD-N cohort. Regions on chromosomes 5 and 17 were suggestive of lin**^2^**kage (p<0.67), with LOD scores of 3.32 and 3.39, respectively. The chromosome 17 locus was later ruled out. The D2.HD-F crosses did not identify linkage. n=25 (19 affected, 6 unaffected^2^); D2=DBA/2J.

#### SJ crosses

Separate QTL analyses of SJ-derived N_2_ and F_2_ recombinants were performed and results are plotted in **Fig. 4** and detailed in Table 2. The SJ.HD-N_2_ recombinants (16 affected and 17 unaffected) found the same distal chromosome 5 QTL identified as a suggestive locus in the D2.HD-N_2_ population, but in this case, the SJ.HD-N_2_ cohort was highly significant (LOD=4.88) and explained 43.2% of the trait variance. With this confirmation, we have designated the chromosome 5 linkage as *Emhd1*, for ‘eosinophilic myocarditis to heart disease’. Interestingly, the SJ.HD-F_2_ population (22 affected and 22 unaffected) did not identify the QTL on chromosome 5. Instead, a highly significant QTL was found on chromosome 17 (Fig. 4B). A closer look at the peak revealed two closely linked loci (Fig. 4C), including a locus proximal to MHC (LOD=7.51; designated *Emhd2*) and another region that contained the MHC region (LOD=6.89; named *Emhd3*). Each locus explained about half of the trait variance. However, their proximity to each other, and the fact that all mice had both QTLs in heterozygosity indicated the variance explained by these QTLs almost entirely overlapped. Similarly, both chromosome 17 QTLs were consistent with dominant variants, such that only one mutant allele of each is required in the HD trait.

**Fig. 4.**
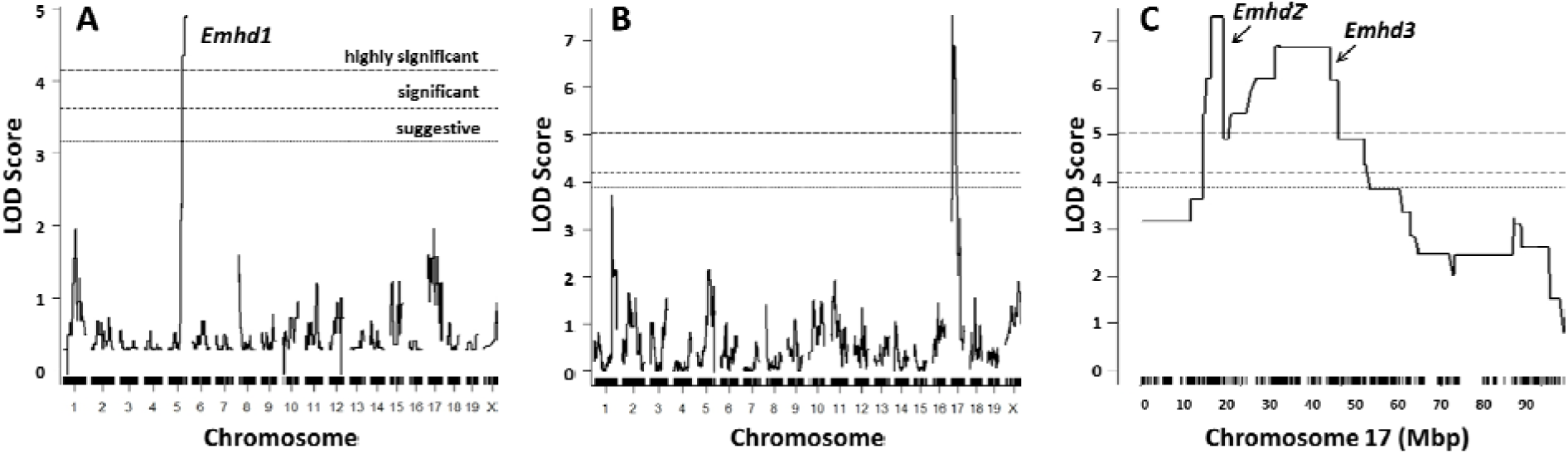
QTL analysis results of the SJ.HD crosses. **A.** SJ.HD-N_2_ (n=33; 16 affected, 17 unaffected); **B.** SJ.HD-F_2_ (n=44; 22 affected, 22 unaffected); and **C.** A closer view of the chromosome 17 peak in B (SJ.HD-F_2_), indicating two distinct QTL peaks. SJ=SJL/J. Suggestive (p<0.67), significant (p<0.05) and highly significant (p<0.01) thresholds for linkage are as indicated in A and determined by 1000 permutations of the respective datasets.

The combined analysis of the 77 SJ-derived N_2_ and F_2_ recombinants (38 affected and 39 unaffected) identified all three *Emhd* peaks (Table 2; Additional file 3). LOD scores were like those from the separate populations, demonstrating the utility of the separate SJ mating schemes to detect recessive versus dominant linkage for this complex trait: the backcrosses identified the recessive *Emhd1* on chromosome 5 and the F_2_ crosses found the dominant-acting *Emhd2* and *Emhd3* on chromosome 17. The height of the *Emhd2* and *Emhd3* peaks flipped in the combined analysis compared with the SJ.HD-F_2_ but were still highly significant. In a 2-QTL model for the chromosome 5 and chromosome 17 (which was treated as one locus) QTLs, the LOD score was additive (13.2), whereas the variance explained (53.6%) remained around half. In addition to the three *Emhd* loci, analysis of this combined SJ.HD-N_2_ + F_2_ dataset also revealed a putative significant linkage on distal chromosome 1 (LOD=4.22; Additional file 3), which was not seen in any of the separate cohorts. However, one can easily see that this peak derives from the smaller peaks at similar positions in the separate analyses (Fig. 4A, 4B); this locus will require further assessment to determine its involvement.

### Haplotype analysis and concordance

Using the Excel file of SNP genotypes for all mice with HD and unaffected littermates used in the mapping cohorts, the heterozygous and both homozygous genotypes were color-coded to visually perform haplotype analysis. This color-coding allowed us to quickly scan the genomewide SNPs, with specific focus on linkage regions, to identify recombinant mice with crossovers that delineated the three QTL intervals. **Fig. 5** presents an overview of the haplotype analysis results. For the recessive *Emhd1* minimal region of effect, heterozygous SJ.HD-F_2_ recombinants determined the proximal and distal ends of the QTL, whereas the bracketed intervals of the dominant *Emhd2* and *Emhd3* QTLs were defined by homozygous SJ alleles in affected SJ.HD-F_2_ recombinants. Backcrosses, which carry at least one copy of the QTLs from the mutant breeder, are necessarily concordant for a dominant mutation (Additional file 4), and thus are uninformative for defining the QTL intervals of *Emhd2* and *Emhd3*. Concordance for dominant loci was therefore assessed in F_2_ mapping cohorts.

**Fig. 5.**
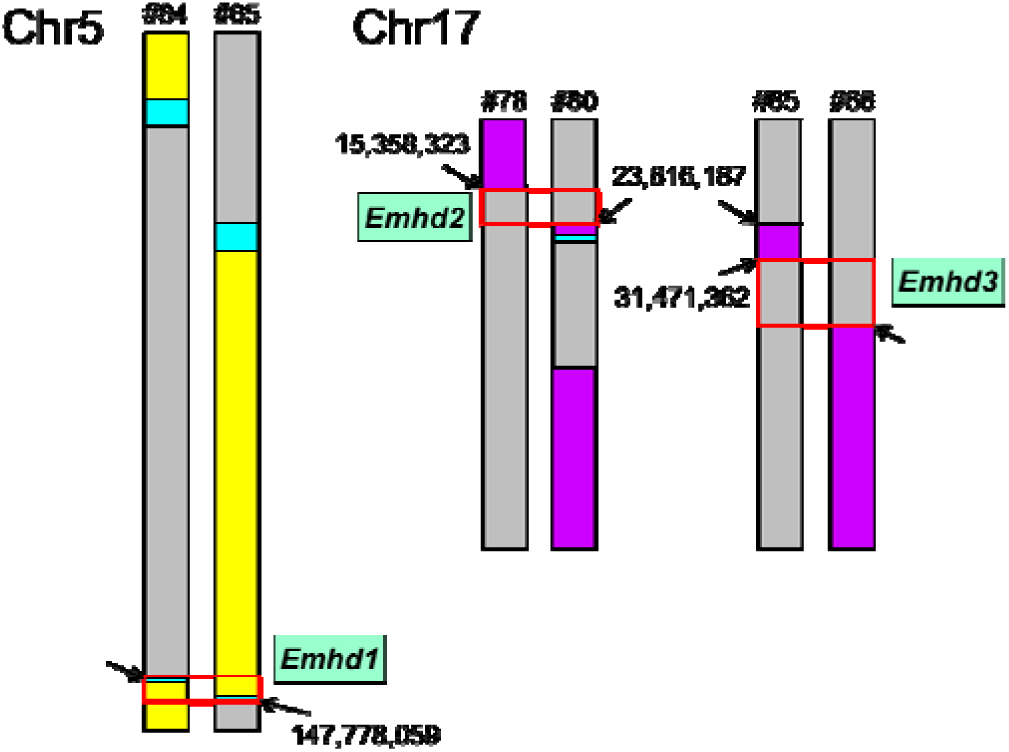
QTL locations and sizes. Haplotype results of the three highly significant *Emhd* QTL intervals (red boxes). Chromosome (Chr) 5 and Chr17 (presented as diploid) of individual numbered recombinants (#) with key crossovers that demarcate the proximal and distal boundaries of each QTL interval. Arrows indicate the SNP positions (GRCm38; Build 38) of each crossover. Yellow=homozygous A/J^HD^; Purple=homozygous SJ; Gray=heterozygous; Cyan=identical by descent.

#### Emhd1

The single B6.HD-N_2_ recombinant (1/50) and the two of 226 S1.HD-F_2_ recombinants that died of HD (Additional file 1) all carried homozygous A/J^HD^-derived genome across *Emhd1*. Among the D2- and SJ-derived recombinant populations, haplotype analysis for *Emhd1* determined large differences in the size of the chromosome 5 interval (Table 2). Specifically, the *Emhd1* interval spanned 22.21 Mb in the SJ.HD-N_2_ population (n=33), which was reduced to 8.58 Mb in the D2.HD-N_2_ cohort (n=25), to 3.57 Mb for the D2.HD-F_2_ recombinants (n=32), and finally down to 2.65 Mb (145.13–147.78 Mb) in the SJ.HD-F_2_ population (n=44). Of note, while neither F_2_ cohort identified linkage to *Emhd1*, individual recombinants of both F_2_ cohorts significantly refined the QTL interval of their respective N_2_ populations. The genotypes of all affected SJ.HD-F_2_ recombinants were concordant with HD across the defined QTL interval (*i.e.,* 22/22 homozygous for mutant A/J^HD^ alleles). However, 3 of 16 D2.HD-F_2_ affected recombinants were discordant (1=heterozygous, and 2=homozygous D2) for the expected homozygous mutant alleles at *Emhd1* (Additional file 4). Among the unaffected SJ recombinant mice in the analysis, *Emhd1* showed the expected 1:1 ratios of heterozygous or homozygous alleles in the N_2_ cohort and ratios of 1:2:1 in the unaffected F_2_ littermates. These data strongly support that *Emhd1* is necessary, but not sufficient to cause EM/HD in this mutant.

#### Emhd2

Haplotype analysis of the D2 crosses for *Emhd2* was not possible, because D2 and A/J inbred strains are identical-by-descent (IBD) for the proximal ∼28.65 Mb of chromosome 17.

However, haplotype analysis of the affected SJ.HD-F_2_ recombinants revealed specific crossovers that defined the proximal and distal ends of *Emhd2*, which confined the *Emhd2* to an 8.46 Mb region (15.36–23.82 Mb; Fig. 5, Table 2) on chromosome 17. This interval maps proximal to the MHC region. All affected SJ.HD-F_2_ mice (22/22) were concordant for a dominant variant (Additional file 4). In contrast, 8/22 unaffected SJ.HD-F_2_ recombinants were discordant (*i.e.,* no mutant alleles), which is consistent with the expectation of ∼1/4 homozygosity (*p*=0.36) for SJ alleles in F_2_ crosses.

#### *Emhd3* and putative QTL on chromosome 1

Haplotype analysis of the 22 SJ.HD-F_2_ affected mice revealed specific crossovers that defined the proximal and distal ends for *Emhd3*. As illustrated in Fig. 5, the *Emhd3* interval was mapped to a 14.19 Mb region (31.47–45.66 Mb; Table 2) on chromosome 17 that includes the major histocompatibility genes and several complement genes of the alternative pathway. All 22 SJ.HD-F_2_ recombinant mice with HD used in the QTL analysis were concordant for a dominant variant (Additional file 4). In addition, the same 8/22 unaffected SJ.HD-F_2_ recombinants that were discordant for *Emhd2* were also discordant for *Emhd3* (*i.e.,* homozygous for SJ alleles throughout both QTL intervals).

Haplotype analysis was also used to determine the interval size of the putative linkage on chromosome 1 (Table 2). Because the genotypes were consistent with a dominant variant (backcrosses are uninformative), only the F_2_ data were used. All 22 SJ.HD-F_2_ affected mice were concordant for a 23.59 Mb interval ranging from ∼171.04–194.63 Mb on distal chromosome 1. Interestingly, the D2.HD-F_2_ cohort was highly discordant (*i.e.,* up to 10/16 affected F_2_ mice were homozygous for D2 alleles across all (6/16) or a large portion (4/16) of this putative chromosome 1 interval, with at least 7/16 affected mice carrying homozygous D2 alleles at every SNP across the interval). This discordance could indicate the chromosome 1 linkage supported by SJ.HD-F_2_ recombinants was a false positive. However, it is also possible that this putative QTL represents a distinction in the set of contributing causal variants for HD between the SJ and D2 crosses.

### Positional candidate genes

Using the Genes and Markers query form in MGI (The Jackson Lab; http://www.informatics.jax.org/marker), we generated lists of all ‘gene feature types’ mapping to the genetic intervals of the three *Emhd* and putative chromosome loci, as determined by haplotype analysis. After removing transcription start sites and CpG island sites, lists of the remaining genetic elements mapping to each of the haplotype-derived QTL intervals were generated, and results were itemized into 8 general categories for reference (Additional file 5). Using literature searches for keywords like heart disease, heart failure, immune dysfunction (including autoimmunity), inflammation, chemokine and cytokine changes, eosinophil pathobiology or coagulopathy, biologically-relevant positional candidates (Table 3) were identified from the known protein coding genes mapping to the highly significant *Emhd* loci. These positional candidate genes represent starting points for follow-on studies.

**Table 3.**
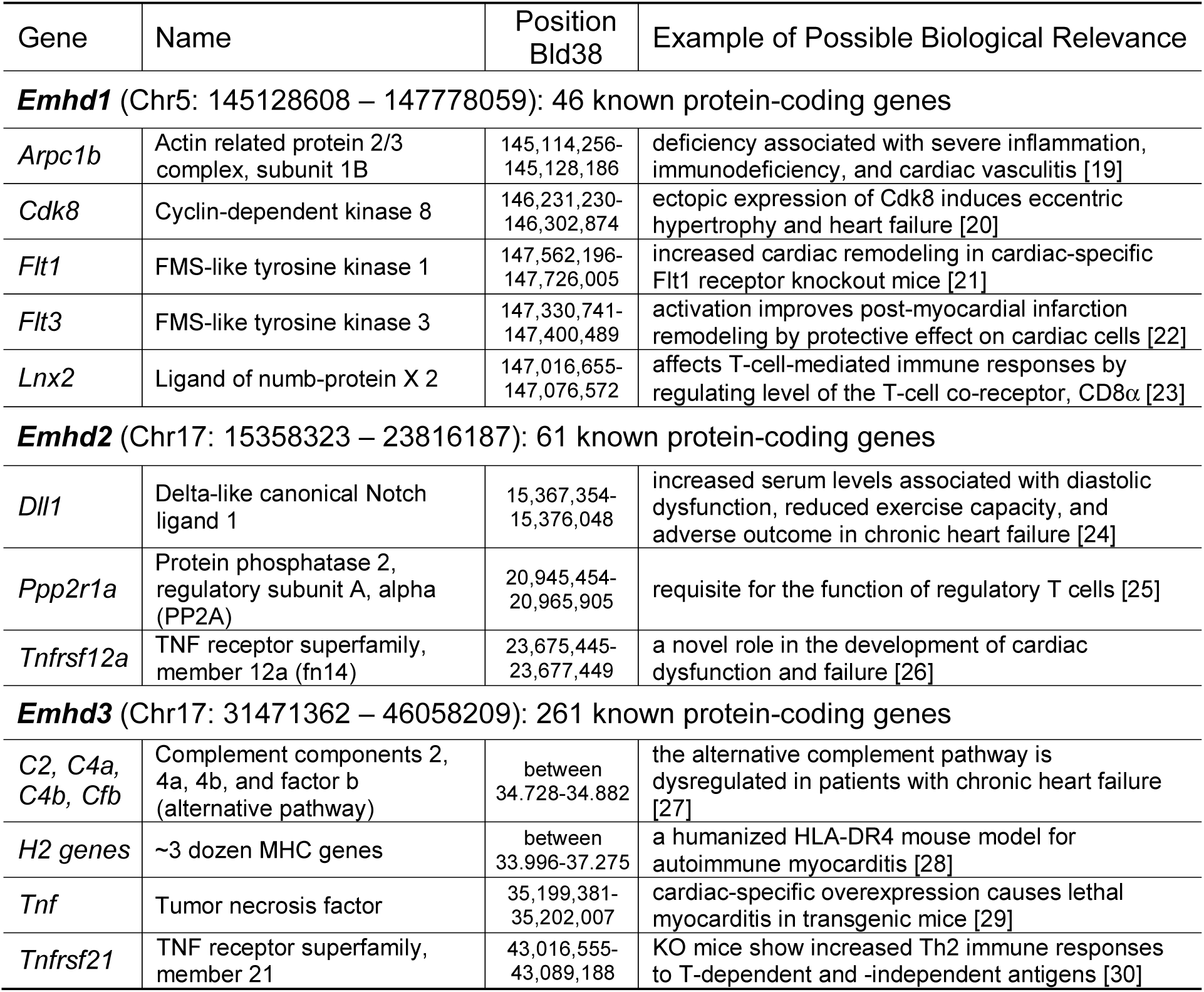
Positional Candidate Genes for *Emhd1-3*.

## Discussion

We have established an inbred mutant line from a founder mouse discovered in the colony. Without a known initiating factor or trigger, affected mice develop EM that progresses to HD and death. We recently reported a detailed histological evaluation of the hearts and lungs of these mutant mice [2]. Although affected mice die with a failing heart, the cardiac pathology was heterogenous. However, these differences can all be ascribed to the spectrum of injuries related to blood and tissue eosinophilia, especially EM and its consequences. In fact, the affected mutant mice present with pathognomonic features of several eosinophil-associated diseases, including natural or induced (*e.g.,* pathogen or autoimmune) EM, hypereosinophilic syndrome (HES) with cardiac complications [31], and eosinophilic granulomatosis with polyangiitis (EGPA; aka Churg-Strauss Syndrome) [32].

Because the eosinophil-associated diseases with heart complications are complex and rare, with few reports demonstrating a familial etiology, little is known regarding the specific genetic determinants involved. Accordingly, genetic analysis for these complex diseases in clinical populations would be difficult, costly, and unlikely to provide the power needed for multigenic discovery. In contrast, the ability to maintain our mutant mouse model of EM/HD for many years strongly supported that the genetic underpinnings of this mutant model were not too complex, and therefore offered an opportunity to map this disease. Two additional findings indicate that HD in this mutant line is controlled by a small set of causal factors. First, we reestablished the original EM/HD trait after several backcrosses and intercrosses to remove the congenic region from the founder; this process recapitulated the disease in an inbred (A/J^HD^) line. Second, mapping studies have revealed three highly significant QTLs, with each locus explaining a significant portion of the trait variance in the relevant crosses. Up to 64% of the variance was explained by a 2-QTL model that included *Emhd1* and either *Emhd2 or Emhd3*, indicating the effects on variance by the two closely mapped chromosome 17 QTLs were not separable in the analysis. These data demonstrate that EM/HD in this mutant line involves the interactions of a limited number of causal factors, for which three highly significant QTLs explain a majority of the trait variance.

To map a disease trait, recombinants from crossing the mutant with another strain must recapitulate all or an essential part of the original trait of interest. Success depends on the number of variants involved in the trait (which is related to penetrance), the mode and direction of inheritance of each mutation, and any non-genetic and/or environmental confounders. The observed rate of affected recombinants depends on the allelic combinations of fixed and segregating causal variants carried between each mating pair. For the A/J^HD^ inbred line, the rates of affected offspring ranged from zero for many breeder pairs to >50% of offspring developing HD for some mating pairs. Based on the variable rates of affected mice over several years of line maintenance, we estimated ∼2-3 variants were likely segregating for the HD trait in the A/J^HD^ line.

To indirectly examine the disconnect between affected rates seen in the A/J^HD^ line versus the few affected mice in the initial three mapping populations, we tested *Dysf* (SJ crosses) and/or *C5* (D2 crosses), two genes with biological relevance to our trait and known to be lacking in A/J mice. DYSF deficiency causes dysferlinopathy [33, 34] and leads to defective membrane resealing in skeletal muscle and to muscle necrosis [35]. DYSF is also involved in cardiomyocyte membrane repair and its deficiency leads to cardiomyopathy in *Dysf* null mice [36, 37]. Therefore, it is plausible that the reduced cardiac repair capacity with *Dysf* deficiency in A/J^HD^ mice contributes to the HD, similar to that associated with other muscular dystrophies [38]. Lack of C5 has also been associated with cardiac failure, but usually after a viral [39], fungal [5, 40] or protozoal [41] infection. Interestingly, it was recently reported that DYSF deficiency leads to increased susceptibility to coxsackievirus B3 (CB3)-induced cardiomyopathy in C5-deficient A/J mice and suggested an important mechanism of DYSF cleavage by viral proteases underpinning cardiac dysfunction [42]. The significant increase in affected mice observed in SJ and D2 crosses indicates the need for direct studies to understand the role for *Dysf* and/or *C5* deficiency in EM/HD.

Many studies have sought to understand the complex genetic control of myocarditis. These studies can be classified by many factors, including spontaneous versus induced; acute versus chronic; type of inducing agent (*e.g.,* pathogen – viral, bacterial or fungal, drug/toxin, or adjuvant); innate, adaptive, and autoimmunity; cellular infiltrate and/or cells affected; and natural animal model versus engineered. Sorting through these studies, we have determined that our mutant mouse model of EM/HD is unique. Prior to our discovery, at least 3 mouse models were known to spontaneously develop EM, including the D2 inbred strain [43], and *Socs1* [44] or *Bcl6* knockout (KO) mice [45]. All other EM models are induced, *e.g.,* cardiac myosin peptides: IL17A^−/−^ IFNγ^−/−^ double knockout [46] or the SWXJ, SJ, or SWR/J mice [47]; or with a pathogen (*e.g.,* CB3 [39, 48]). But, unlike D2 mice that resolve the spontaneous EM by ∼8-10 weeks old [49], the naturally-occurring EM of our mutant line rapidly progresses through myocardial damage, inflammation and thrombotic events that can cause sudden cardiac death or progress to DCM and fatal HD. These outcomes differed from other mouse models of spontaneous EM [43–45] and parallel the more egregious pathologies in HES and EGPA patients. Interestingly, a combined KO-transgenic mouse model generated with human HLA receptor (*e.g.,* HLA DQ8.NOD Aβo) spontaneously develops myocarditis and DCM [50–52], but this myocarditis was not related to eosinophil damage. Thus, in addition to its use as a genetic tool, our mouse model is unique for presenting a spontaneous fatal EM due to cardiac manifestations. As such, the mutant line also provides a means to compare with other models of myocarditis to understand differentiating biological and pathological roles of the eosinophil in the progression from myocarditis to HD, and to understand differences in disease initiation (natural/induced) between models.

Lending validity to our QTL findings, the two putative QTLs we identified on chromosome 17 coincide with previously reported loci in a CB3-induced mouse model of chronic autoimmune myocarditis [53]. In that report, the B6.A-17 chromosome substitution strain [54] was found to confer susceptibility to CB3-induce myocarditis. The genetic background of B6.A-Chr17 mice is inbred B6, except for the A/J-derived chromosome 17. These B6.A-Chr17 consomic mice were bred to produce various chromosome 17 congenic lines carrying different portions of chromosome 17 to use as recipient strains for viral-induced myocarditis. The study identified four regions on chromosome 17 harboring susceptibility loci for CB3-induced myocarditis, including an MHC locus and 3 putative loci proximal or distal to the MHC complex. The distal locus on chromosome 17 was ruled out in our mutant model, because many recombinants with EM/HD carried homozygous SJ alleles across that region. The other three loci could coincide with the *Emhd2* (upstream of MHC genes) and *Emhd3* (this large region may house more than one QTL) intervals. Two differences between these models are clear: (1) our model of myocarditis, as far as we know, does not require induction and (2) the EM/HD mutant primarily involves an eosinophilic infiltration that causes cardiac inflammation, whereas the CB3-induced myocarditis is characterized by a diffuse mononuclear interstitial infiltrate [55]. We suspect that the non-chromosome 17 locus *Emhd1* on chromosome 5 could help explain at least some of the difference, especially regarding the role of eosinophils.

Two limitations for these studies also need mentioning. First, as stated above, D2 mice spontaneously develop a non-lethal course of EM at a very young age, which usually resolves by 10-weeks old [43]. Therefore, it is possible that some of the D2-related recombinants dying of HD were confounded by this early-age EM and the susceptibility alleles gained in the crosses with A/J^HD^. This could also explain why the D2 data was less effective at identifying significant linkage. Similarly, this confounder could be the reason why 3/16 D2.HD-F_2_ mice were discordant at *Emhd1*, whereas all other N_2_ and F_2_ mice with HD carried homozygous A/J^HD^ mutant alleles (Additional file 4). One could speculate that the *Emhd1* causal variant resides in a gene or pathway involved in the resolution of EM in D2 mice. Second, the ‘unaffected mice’ used in these studies were littermates that did not show any signs of illness at sacrifice (∼20-weeks old). However, we had several breeders used to generate the recombinant populations that died of HD much later than 140 days old. Therefore, the ‘unaffected’ designation does not account for the possibility that they would have developed HD later.

## Conclusion

Using crosses of four inbred strains with A/J^HD^ mutant males we generated eight different recombinant cohorts of N_2_ and F_2_ mice to recapitulate the EM/HD trait for linkage analysis. This methodical progression dramatically improved efficiency to produce offspring with HD from 0 to >17%, which was critical for mapping this oligogenic trait. QTL analysis of cohorts of affected recombinants and siblings with no HD by 20-weeks old revealed three highly significant QTLs that can explain nearly two-thirds of the genetic variance. The consistent >2-fold increase in affected D2 and SJ recombinants, suggests *Dysf* (or another SJ variant, or SJ pathology in common with A/J) and lack of C5 (or other D2 change in common with A/J) may also have unknown modifying roles in susceptibility to EM/HD and could help explain the remaining missing heritability. As few genes for eosinophil-related diseases are known in mouse or man, finding the causal genes that can explain this mouse model of EM/HD would allow directed studies to reveal pathological mechanisms and understand disease history, with a goal to interrogate potential translation to relevant human diseases.

## Methods

### Mice

All inbred mouse strains were obtained from the Jackson Laboratory (JAX: Bar Harbor, ME) and all lines and crosses generated and used in these studies were derived from the original inbred strains. Mice were housed within a specific pathogen free AALAC approved facility at Cincinnati Children’s Hospital. Mice were placed in shoebox containers with corncob bedding at up to 4 animals per cage, and a 12-hour on/12-hour off light/dark cycle. Animals were checked daily by Veterinary Services, including routine husbandry and cage changes, as well as daily status checks by a Compliance Technician. Laboratory staff performed routine monitoring of the colony at least 3 times a week, including necessary matings, weanings, ear tagging, phenotype observation, and animal sacrificing, as needed. Mice found to be ungroomed, immobile, dyspnic, cyanotic, or showing other signs of distress or illness will be used or sacrificed promptly, usually that day. When identified by Veterinary Services staff, they send a text message directly to the PI and an e-mail to staff, alerting us of a ‘sick’ mouse.

Because mice usually do not show signs of illness until late in disease, usage of sick mice is addressed at that time – mice are sacrificed by Veterinary Services staff or, if needed by the lab, taken to necropsy for immediate use or sacrifice to obtain biological specimens. On weekends, the PI is called for instructions on how to handle the sick mouse (usually sacrificed by Veterinary Services staff). Moribund mice that were sacrificed and all mice found dead were visually assessed to verify HD. Recombinant mice were deemed ‘unaffected’ if they survived to 140-days old (20 weeks) without signs of disease, a time when most affected mutant mice developed HD. All experimental mice used in these studies, including unaffected mice, were sacrificed by CO_2_ or isoflurane inhalation overdose before sample collections (*e.g.,* tail clip, blood, tissue, and organ samples collected, as needed). All mice found moribund were used immediately or sacrificed by CO_2_ inhalation. Protocols were reviewed and approved by the Cincinnati Children’s Hospital Institutional Animal Care and Use Committee (IACUC protocol #2016-0048 to DRP), which required annual renewals and a total rewrite every three years.

### Discovery of the mutant line

The founder mutant mouse was initially found in an A.B6-Chr17 congenic line already in the colony for another purpose. The genetic background of the A.B6-Chr17 congenic was inbred A/J, except for a ∼45-Mb region of C57BL/6J (B6) introgressed onto distal chromosome 17 (as verified by the MegaMUGA SNP panel; GeneSeek, Neogen Genomics; Lincoln, NE). To determine whether the chromosome 17 congenic B6 interval contributed to the mutant trait, the B6 congenic region was removed by backcrossing with inbred A/J mice (JAX stock #000646), intercrossing first-generation siblings (A/J with a heterozygous congenic interval) and genotyping the second-generation offspring for microsatellite markers (IDT; Coralville, IA) polymorphic for A/J and B6 that spanned the congenic interval. Following the removal of the B6 congenic interval, the A/J inbred status of the mutant line was again verified using the Illumina-based MegaMUGA panel of ∼77,800 SNPs (GeneSeek). Importantly, after reestablishing the A/J inbred background, we were able to recapitulate EM/HD (A/J^HD^), but only after several breeder pairs and multiple litters from each pairing, suggesting more than one gene was still segregating for the trait. Generating mice that developed EM/HD after the congenic interval was removed verified that the B6 interval (and the A/J complement region that replaced it) was not contributing to the mutant trait. To help ensure that we retain the causal alleles in the gene pool, numerous sibling-sibling or offspring-parent mating pairs are maintained at each generation to propagate the trait. Successfully reestablishing EM/HD in the inbred line not only demonstrated trait heritability but suggested that its mode of inheritance was not overly complicated.

### Phenotype

The full characterization of the mutant A/J^HD^ line has recently been described [2]. In that report, the mutant line is referred to as AJ.EM, since EM was the primary early histological finding. The diseased heart develops a profuse EM and inflammatory dilated cardiomyopathy (DCM), often with considerable right ventricular outer wall expansion to a point of translucency. Some mutant hearts demonstrated pericardial fibrosis and less severe HD, with an underlying atrial or ventricular thrombus that likely caused sudden death. As seen in the inbred A/J^HD^ mutant line, affected recombinant mice from the various crosses also exhibited gross signs of HD at or near death, including subcutaneous edema, cardiomegaly, and fibrosis. Accordingly, the presence of HD was used as the trait for mapping in these studies, since it was universal in affected mice and easy to verify visually at necropsy.

### Whole mounts and Trichrome staining

The hearts from siblings of the A/J^HD^ mutant line were processed for whole-mount imaging to demonstrate size and structural variability among affected and unaffected mice of the litter. Hearts from littermates were taken when the first sibling showed signs of HD (*e.g.,* panting, inactivity, ungroomed, and weight gain from pericardial and peripheral edema). Hearts were excised, cleared in phosphate-buffered saline, photographed with a dissecting microscope at low setting, coronally cut through the 4 chambers and re-photographed at the same settings. Hearts were then placed in 10% phosphate-buffered formalin overnight and submitted to the Cincinnati Children’s Research Pathology Core for standard processing, paraffin embedding and staining with Trichrome.

### Mapping studies

Females from four different inbred strains were mated with proven A/J^HD^ mutant males (*i.e.,* previously sired at least one offspring with verified HD) to generate recombinants for mapping the EM/HD trait. All inbred strains used in crosses were originally obtained from The Jackson Laboratory (JAX; Bar Harbor, ME) and maintained in-house as small cohorts for breeding needs. If maintained in the colony, inbred strain breeders were replaced after five generations to preserve genetic background integrity with the ‘Jackson’ substrain. Inbred mice used for QTL mapping included: 129S1/SvImJ (S1; JAX stock #002448); C57BL/6J (B6; JAX stock #000664), DBA/2J (D2; JAX stock #000671), and SJL/J (SJ; JAX stock #000686). Populations for mapping the HD trait were generated using backcross (N_2_) and/or intercross (F_2_) breeding schemes among these four inbred strains (Additional file 6). Recombinant mice from all crosses were closely monitored and aged up to 20-weeks old to identify mice with HD, an age that accounted for ∼85% of all deaths in affected mice of the A/J^HD^ line. The affected populations for QTL analysis also included a few older breeders with verified HD. Unaffected recombinant mice used in QTL analysis included littermates without signs of HD (*e.g.,* behavior and appearance; heart at necropsy) at sacrifice (at least 20-weeks old).

### Genotyping

To remove the B6 congenic region on chromosome 17 in the original founder mutant, microsatellite markers (IDT; Coralville, IA) polymorphic between A/J and B6 mice were typed, including *D17Mit178* (48.71 Mb; Bld38), *D17Mit139* (52.66 Mb), *D17Mit88* (57.37 Mb), and *D17Mit39* (74.28 Mb). DNA isolation, PCR and gel electrophoresis were performed as described previously [56, 57]. For linkage analysis, DNA isolated from tail tips of affected and unaffected mice from the A/J^HD^ crosses were genotyped for ∼143,000 SNPs using the GigaMUGA array [58] at GeneSeek (Lincoln, NE; a subsidiary of Neogen Corporation). A total of 134 DNA samples with high-quality SNP results were used in the QTL analyses.

### QTL and haplotype analyses

Linkage analysis was performed using the R/qtl software package [16, 17], after checking quality and intensity normalization of the SNP data using routines provided in the R-package argyle [18]. Affected and unaffected N_2_ and F_2_ recombinants from D2- or SJ-derived crosses were analyzed separately to identify cross-specific QTLs. For linkage analysis, the GigaMUGA SNP profile was reduced to the informative markers and the trait designated as the presence or absence of HD. Genomewide empirical thresholds were obtained within the R/qtl program. Following QTL analysis, the whole genome (with specific focus on the linkage regions) was inspected by haplotype analysis using genotype color-coding in an Excel spreadsheet. Haplotype analysis allowed us to identify specific recombinant mice with crossovers that defined the minimal QTL interval associated with the HD trait and to predict the likely mode of inheritance of each locus.

### Statistical analysis

Differences in HD rates between the sexes (2 categorical variables) were assessed for each cross using a Chi-square test and *p*<0.05 for significance. QTL significance levels for each analysis were determined using at least 1,000 permutations of the respective datasets [59]. Genomewide threshold levels for linkage were set at *p*<0.01 (highly significant), *p*<0.05 (significant) and *p*<0.67 (suggestive).

## Abbreviations

A/J: inbred mouse strain A from Jackson Lab (J)
B6: C57BL/6J inbred mouse strain
C5: hemolytic complement
CB3: Coxsackievirus B3
Chr: chromosome
Emhd: quantitative trait, eosinophilic myocarditis to heart disease
D2: inbred mouse strains DBA/2J
Dysf: dysferlin protein (DYSF) or gene (*Dysf*)
EGPA: Eosinophilic granulomatosis with polyarteritis (aka Churg-Strauss Syndrome)
EM: eosinophilic myocarditis
HD: heart disease
HES: Hypereosinophilic Syndrome
IBD: identical-by-descent
JAX: Jackson Laboratory
LOD: a logarithmic score representing the likelihood of linkage, specifically the log of the <u>od</u>ds ratio for linkage to non-linkage
Mb: megabases
MUGA: mouse universal genotyping array
QTL: quantitative trait locus
SJ: mouse inbred strain SJL/J
SNP: single nucleotide polymorphism

## Declarations

## Acknowledgments

We thank Ms. Hanna Osinska, Ph.D. for whole-mount imaging and coronal cuts of the control and mutant mouse hearts and the serial Trichrome imaging. We acknowledge the many dedicated laboratory staff and students over the years that managed and closely monitored the mutant mouse colony.

## Availability of data and materials

The datasets used and/or analyzed during the current study are available from the corresponding author on reasonable request.

## Author’s contributions

WJG and SMH maintained the mouse lines, managed all the crosses, genotyped and phenotyped the mice, isolated DNA, and performed studies. WJG also performed the scanone and scantwo QTL analyses and generated associated QTL plots. NZ provided expertise and advise in pathology, interpretations of histopathology slides, and helped write, edit and review the manuscript. DRP conceived of the studies and oversaw the project, analyzed the data, generated all tables and final figures, and wrote the original draft of the manuscript. All coauthors read, provided input and approved submission of the manuscript.

## Ethics approval

Mice were handled in accordance with the protocols approved by the Institutional Animal Care and Use Committee at Cincinnati Children’s Hospital Medical Center Protocol #: IACUC 2016-0048 (DRP).

## Consent for publication

Not applicable.

## List of additional files

**Additional file 1.** File contains a figure that shows an overview and outcomes from 4 rounds of breeding crosses to map the EM/HD trait (pptx 231 kb).

**Additional file 2.** File contains the QTL analysis results of D2.HD-F_2_ and the combined D2.HD-N_2_ and D2.HD-F_2_ cohorts (pptx 224 kb).

**Additional file 3.** File contains the QTL analysis results of the combined SJ.HD-N_2_ and SJ.HD-F_2_ cohorts (pptx 195 kb).

**Additional file 4.** File contains a table for the concordance of QTL genotypes with HD in recombinant mice used in the QTL analysis (docx 37 kb).

**Additional file 5.** File contains a table listing the positional candidate gene elements for the three QTLs (docx 37 kb).

**Additional file 6.** File contains a figure summarizing the breeding schemes for mapping the HD trait (pptx 83 kb).

