## Additional File 1. for "Heart Disease in a Mutant Mouse Model of Spontaneous Eosinophilic Myocarditis Maps to Three Highly Significant Loci"

### Slide 1
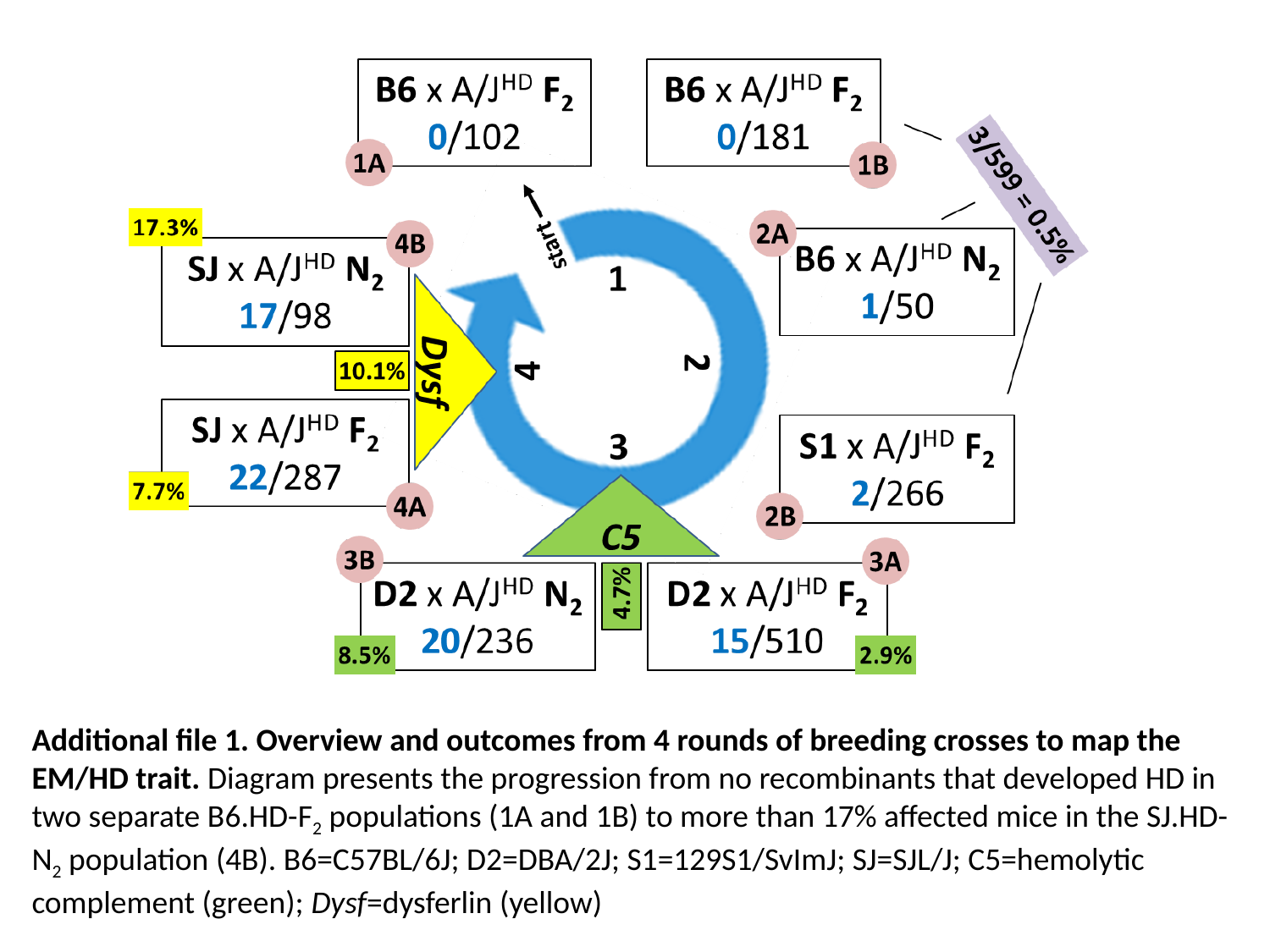

Additional file 1. Overview and outcomes from 4 rounds of breeding crosses to map the EM/HD trait. Diagram presents the progression from no recombinants that developed HD in two separate B6.HD-F_2_ populations (1A and 1B) to more than 17% affected mice in the SJ.HD-N_2_ population (4B). B6=C57BL/6J; D2=DBA/2J; S1=129S1/SvImJ; SJ=SJL/J; C5=hemolytic complement (green); Dysf=dysferlin (yellow)
