## Additional File 2. for "Heart Disease in a Mutant Mouse Model of Spontaneous Eosinophilic Myocarditis Maps to Three Highly Significant Loci"

### Slide 1
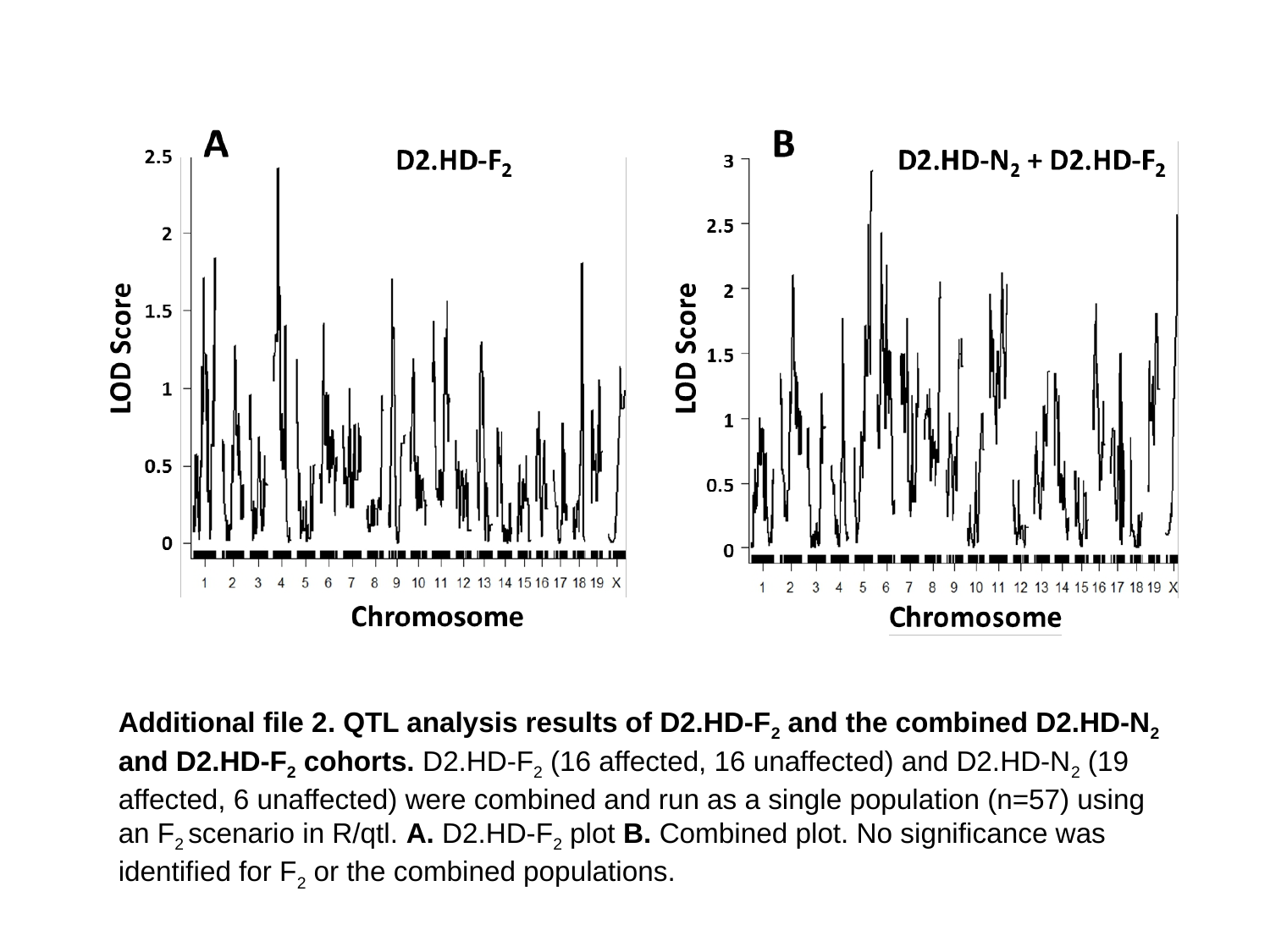

Additional file 2. QTL analysis results of D2.HD-F_2_ and the combined D2.HD-N_2_ and D2.HD-F_2_ cohorts. D2.HD-F_2_ (16 affected, 16 unaffected) and D2.HD-N_2_ (19 affected, 6 unaffected) were combined and run as a single population (n=57) using an F_2_ scenario in R/qtl. A. D2.HD-F_2_ plot B. Combined plot. No significance was identified for F_2_ or the combined populations.
