## Additional File 3. for "Heart Disease in a Mutant Mouse Model of Spontaneous Eosinophilic Myocarditis Maps to Three Highly Significant Loci"

### Slide 1
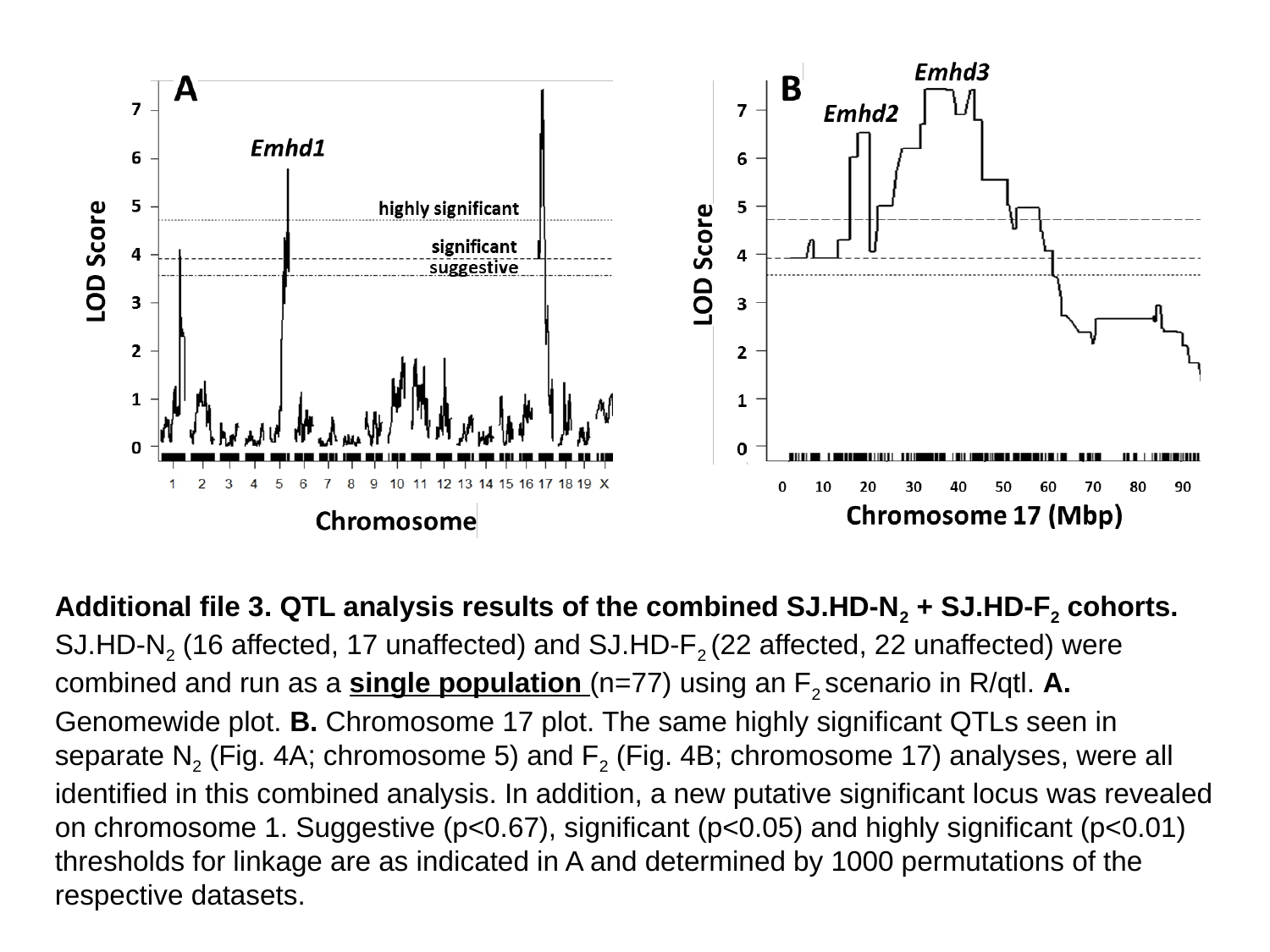

Additional file 3. QTL analysis results of the combined SJ.HD-N_2_ + SJ.HD-F_2_ cohorts. SJ.HD-N_2_ (16 affected, 17 unaffected) and SJ.HD-F_2_ (22 affected, 22 unaffected) were combined and run as a single population (n=77) using an F_2_ scenario in R/qtl. A. Genomewide plot. B. Chromosome 17 plot. The same highly significant QTLs seen in separate N_2_ (Fig. 4A; chromosome 5) and F_2_ (Fig. 4B; chromosome 17) analyses, were all identified in this combined analysis. In addition, a new putative significant locus was revealed on chromosome 1. Suggestive (p<0.67), significant (p<0.05) and highly significant (p<0.01) thresholds for linkage are as indicated in A and determined by 1000 permutations of the respective datasets.
