## Additional File 4. for "Heart Disease in a Mutant Mouse Model of Spontaneous Eosinophilic Myocarditis Maps to Three Highly Significant Loci"

**Additional file 4.** Concordance of QTL genotypes with HD in recombinant mice.

|  | **Recombinant Cross** | | | |
| --- | --- | --- | --- | --- |
| **Linkage** | D2.HD-N_2_ | D2.HD-F_2_ | SJ.HD-N_2_ | SJ.HD-F_2_ |
| ***Emhd1***  (5:145128608-147778059) | 20/20 | 13/16 | 16/16 | 22/22 |
| ***Emhd2***  (17:15358323–23816187) | **IBD *** | IBD * | **16/16** | 22/22 |
| ***Emhd3***  (17:31471362–45658739) | **20/20** | 13/16 | **16/16** | 22/22 |
| 1:171038979 – 194625219  (significant QTL from combined SJ crosses) | **20/20** | 6/16 ^#^ | **16/16** | 22/22 ^#^ |

*Emhd1* correlates with a recessive variant (2 mutant alleles); *Emhd2* and *Emhd3* correlate with dominant variants (one mutant allele).

* The proximal ~28.65 Mb of chromosome 17 is identical-by-descent (IBD) between DBA/2J (D2) and A/J (and A/J^HD^) strains, so information on allelic origin cannot be obtained.

^#^ Putative QTL on chromosome 1 and haplotype concordance was found only in SJ.HD-F_2_ cohort.

Cells with **bold** text represent backcrosses that are necessarily concordant for a dominant trait, as they always carry at least one mutant allele.
