## Additional File 5. for "Heart Disease in a Mutant Mouse Model of Spontaneous Eosinophilic Myocarditis Maps to Three Highly Significant Loci"

**Additional file 5. Positional candidate gene elements.**

| Gene feature | *Emhd1*  5:145128608–147778059 | *Emhd2*  17:15358323–23816187 | *Emhd3*  17:31471362–45658739 | Distal Chr1  1:171038979–194625219 |
| --- | --- | --- | --- | --- |
| protein coding | 46 | 107 | 336 | 236 |
| lncRNA | 23 | 36 | 109 | 200 |
| QTL | 4 | 18 | 50 | 95 |
| snoRNA | 3 | 1 | 17 | 22 |
| tRNA / rRNA | 0 / 0 | 8 / 0 | 0 / 0 | 29 / 2 |
| miRNA | 1 | 5 | 19 | 19 |
| pseudogene | 27 | 114 | 181 | 99 |
| unclassified  gene / ncRNA | 6 / 0 | 5 / 0 | 16 / 12 | 70 / 9 |
| Total | 110 | 294 | 740 | 781 |
