## Additional File 6. for "Heart Disease in a Mutant Mouse Model of Spontaneous Eosinophilic Myocarditis Maps to Three Highly Significant Loci"

### Slide 1
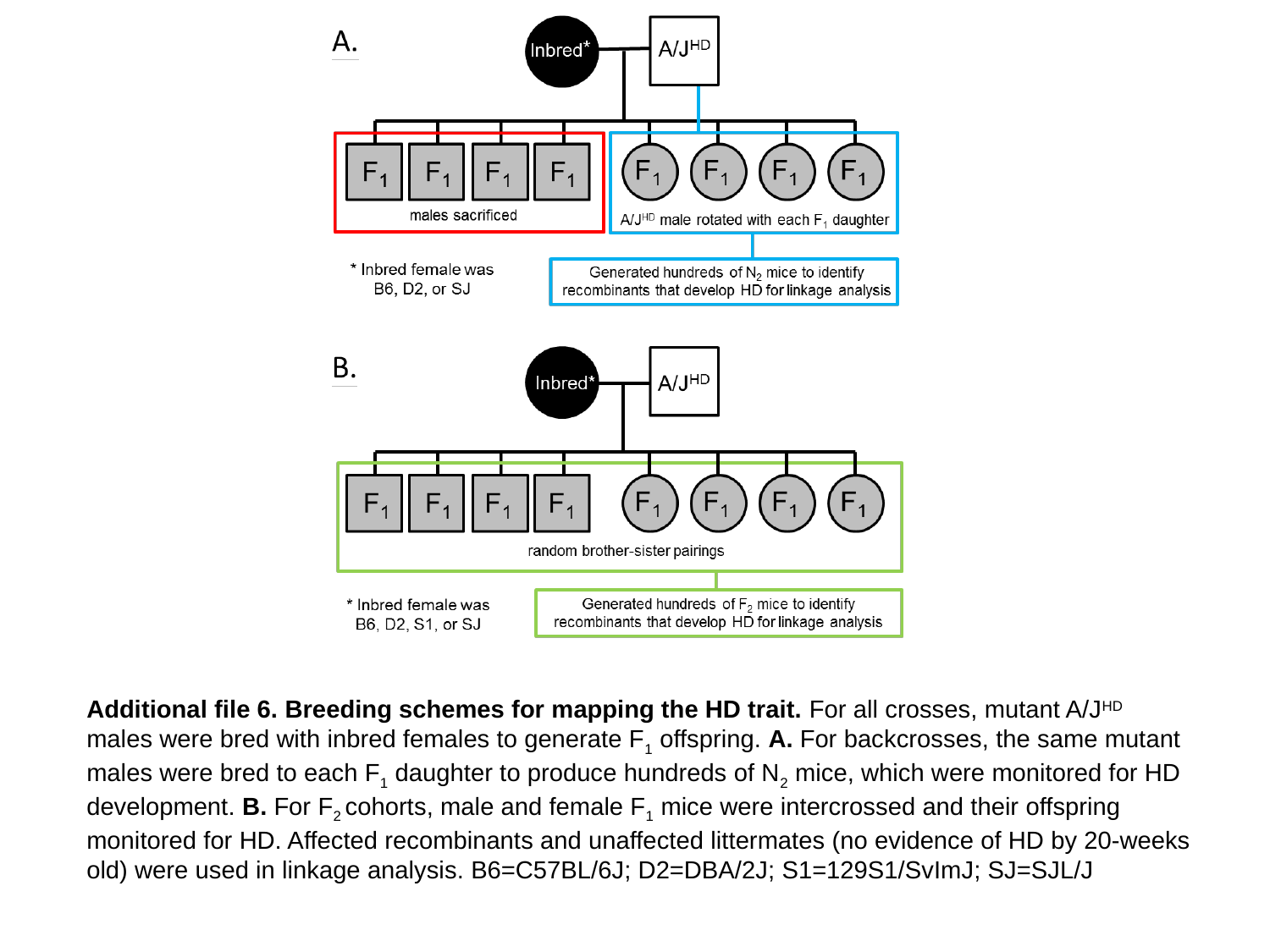

Additional file 6. Breeding schemes for mapping the HD trait. For all crosses, mutant A/J^HD^ males were bred with inbred females to generate F_1_ offspring. A. For backcrosses, the same mutant males were bred to each F_1_ daughter to produce hundreds of N_2_ mice, which were monitored for HD development. B. For F_2_ cohorts, male and female F_1_ mice were intercrossed and their offspring monitored for HD. Affected recombinants and unaffected littermates (no evidence of HD by 20-weeks old) were used in linkage analysis. B6=C57BL/6J; D2=DBA/2J; S1=129S1/SvImJ; SJ=SJL/J
